# Distinct motivations to seek out information in healthy and addicted individuals

**DOI:** 10.1101/2020.02.19.955989

**Authors:** Irene Cogliati Dezza, Xavier Noel, Axel Cleeremans, Angela J. Yu

**Affiliations:** Centre for Research in Cognition & Neurosciences, ULB Neuroscience Institute, Université Libre de Bruxelles, Belgium; Department of Experimental Psychology, Faculty of Brain Sciences, University College London, London, UK; Faculty of Medicine, Université Libre de Bruxelles, Belgium; Department of Cognitive Science, University of California San Diego, United States

**Author notes:** **Supplemental Information:** Yes.

**Keywords:** Novelty-seeking, Information-seeking, Exploration, Gambling Disorder, Reinforcement learning

## Abstract

As massive amounts of information are becoming available to people understanding the mechanisms underlying information-seeking is more pertinent today than ever. In this study, we investigate the underlying motivations to seek out information in healthy and addicted individuals. We developed a novel decision-making task and a novel computational model which allow to dissociate the relative contribution of two motivating factors to seek out information: a desire for novelty and a desire to reduce uncertainty. To investigate whether/how the motivations to seek out information vary between healthy and addicted individuals, in addition to healthy controls we included a sample of individuals with gambling disorder- a form of addiction without the confound of substance consumption and characterized by compulsive gambling. Our results indicate that healthy subjects and problem gamblers adopt distinct information-seeking “modes”. Healthy information-seeking behavior was mostly motivated by a desire for novelty. Problem gamblers, on the contrary, displayed reduced novelty-seeking and an increased desire to reduce uncertainty (general information-seeking) compared to healthy controls. Our findings not only shed new light on the motivations driving healthy and addicted individuals to seek out information, but they also have important implications for treatment and diagnosis of behavioral addiction.

## INTRODUCTION

Recent advancements in neuroscience have shown information-seeking to be an essential aspect of human cognition that supports healthy decision-making and goal-directed processing ^1 2 3 4 5 6 7 8^. Information-seeking is often contraposed to the human tendency of maximizing immediate benefits (reward-seeking). A decision-maker who is trying to find out the best restaurant in town may try out all different available options in order to obtain information on the potential benefit of each restaurant, but this information search may be costly or result in unpleasant experiences.

Yet, healthy humans finely balance the urge for immediate reward vs. longer-term information gain during repeated choice behavior, thus negotiating an exploration-exploitation trade-off ^4 6^; ^9^. On the contrary, in certain psychopathologies such as addictive behaviors resolving this tension is highly compromised resulting in reduced information-seeking ^10^. Previous studies have suggested that a desire for *novelty* (^11 12 1 13 14 15^ and a drive to seek out *general information* ^16 17^, may both drive human information-seeking behavior. However, there has been no study that systematically analyzes the relative importance of these two factors in healthy humans, nor how this information-seeking system might be altered in addictive behaviors.

While novelty is only associated with a completely novel item, uncertainty-reduction can promote the exploration of an option beyond the first encounter. These two motivational factors are however highly related since the uncertainty/information bonus is highest for a novel option. Thus, an uncertainty/information bonus and a novelty bonus can be easily mistaken for one another as statistically significant explanatory factors. However, these two motivational factors seem to rely on different neural regions in the brain, with novelty-seeking expressed in midbrain dopaminergic regions ^18 11 12 19^ and general information seeking in prefrontal regions ^20 21 22^. Here, we state that the distinction between novelty-seeking and general information-seeking is essential to understand the underlying motivations to seek out information in healthy and addicted individuals.

For example, evidence for general information-seeking has come from variants of sequential learning and decision-making tasks (e.g., the bandit tasks; ^4 6 8 23^. This may leave the possibility that general information-seeking is more important for scenarios in which repeated choices are necessary such as during learning or planning, while novelty-seeking might be more relevant for single-stage decisions or early stages of learning ^12 14^. Additionally, impaired information-seeking in addictive disorders ^10 24^ has been explained as a general reduction in the desire to reduce uncertainty about the environment. However, these impairments might be equally explained as a reduced desire for exploring novel opportunities or engaging in novel behavioral patterns. This distinction is crucial for addictive behaviors - of which gambling disorder is a prototype ^25^. If reduced information-seeking is caused by a reduced motivation specifically for exploring novel options, this could explain why pathological gamblers exhibit perseverance in behavioral routines despite the negative consequences associated with them (e.g., financial loss^25^, but at the same time they still prefer choices associated with high uncertainty about reward outcome (e.g., gambling games such as gaming machines or blackjack ^26 27^. Insight into the distinction between novelty-seeking and general information-seeking is therefore particularly relevant for understanding addictive disorders, as well as potentially developing better diagnostic tools or clinical treatments.

Here, we explicitly compare general information-seeking and novelty-seeking in a modified version of the bandit task, which makes it possible to dissociate the relative contribution of expected reward, novelty, and general information as motivating factors in choice behavior. We also implement a reinforcement-learning type model to quantitatively separate out the importance of these three factors in driving human choice behavior. In addition to healthy controls (HCs), we include a sample of individuals with gambling disorder (PGs). This allows us to investigate the relative contribution of general information-seeking and novelty-seeking in addictive behaviors.

## METHODS AND MATERIAL

### Participants

Forty (40) unmedicated PGs (mean age = 30.1, 4 females) and twenty-two (22) HCs (mean age = 29.0, 4 females) were recruited from the local communities (**Table 1**; **Supplement**). The sample size of both groups was based on previous studies ^28 6^. Gamblers were selected among those who were gambling at least once per week, while HCs were those without gambling experience in the year preceding experimental participation (**Table 1**; **Supplement**).

**Table 1.** Demographic information. Mean and standard deviations are shown for each measure. For each comparison, we ran a two-sampled t test, except for gender comparison where chi-squared test was used. The two groups differ only in terms of gambling severity (with no gambling problems reported in the control group) and years of education as often reported in the literature ^28^ (years of education did not correlate with any of the behavioral measures considered in this study and removing PGs with lower years of education did not change the main results reported in the text). Note: WAIS IV-Wechsler Adult Intelligence Scale (the block-design component of the WAIS is the subset that best predicts performance IQ ^29^); CPGI-Canadian Problem Gambling Index; AUDIT - Alcohol Use Disorders Identification Test; DAST - Drug Abuse Screening Test; FTND - Fagerström Test for Nicotine Dependence; ACS - Attentional Control Scale; BDI- Beck Depression Inventory; STAI-S - State version of the State-Trait Anxiety Inventory; STAI-T - Trait version of the State-Trait

|  | PGs<br>n=40 | HCS<br>n=22 | Test Statistic |
| --- | --- | --- | --- |
| Gender (M/F) | 36\|4 | 18\|4 | $p = 0.601$ |
| Age | 30.1(9.3) | 29.0(6.6) | $p = 0.982$ |
| Years of Education | 14.7(2) | 16.2(2.2) | $p = 0.037^*$ |
| IQ (WAIS block) | 8.4(2.6) | 9.3(1.9) | $p = 0.131$ |
| Gambling Severity (CPGI) | 8.8(6.1) | 0 | $p < 10^{-10}^*$ |
| Alcohol use (AUDIT) | 4.6(3.9) | 5.3(3.1) | $p = 0.48$ |
| Drug use (DAST) | 0.225(0.423) | 0.227(0.429) | $p = 0.992$ |
| Smoking dependence (FTND) | n=4 | n=1 | NA |
| Memory Capacity (WAIS) | 10.3(3.5) | 9.7(4.1) | $p = 0.483$ |
| Attentional Control (ACS) | 35.4(9) | 37.5(7) | $p = 0.312$ |
| Depression (BDI) | 5.6(4.9) | 4.2(4.8) | $p = 0.137$ |
| Anxiety (STAI-S) | 35.1(10.9) | 37.9(9.5) | $p = 0.173$ |
| Anxiety (STAI-T) | 39.6(12.4) | 43.1(11) | $p = 0.2$ |
| Positive Mood (PANAS) | 35.4(6.3) | 36.3(5.3) | $p = 0.701$ |
| Negative Mood (PANAS) | 21.1(7.9) | 19.8(4.8) | $p = 0.808$ |

### Behavioral Task

Participants performed 162 games of a decision-making task ^6^, which permits the dissociation of reward and information on sequential choices ^4^ **Figure 1a, Supplement**). Each game consists of two phases (or tasks): participants were initially instructed about which option (deck of cards) to choose from on each trial (*forced-choice task*; **Figure 1b**) for six consecutive trials, after which they were free to choose from any of the options (*free-choice task*; **Figure 1c**) so as to maximize their total gain. The number of free-choice trials varied from 1 to 6 trials, and was inverse-exponentially distributed, such that subjects were most frequently allowed to make 6 free choices. The total gain was shown to the subject at the end of the experiment and converted to a monetary payoff (0.01 euros for every 60 points).

**Figure 1.**
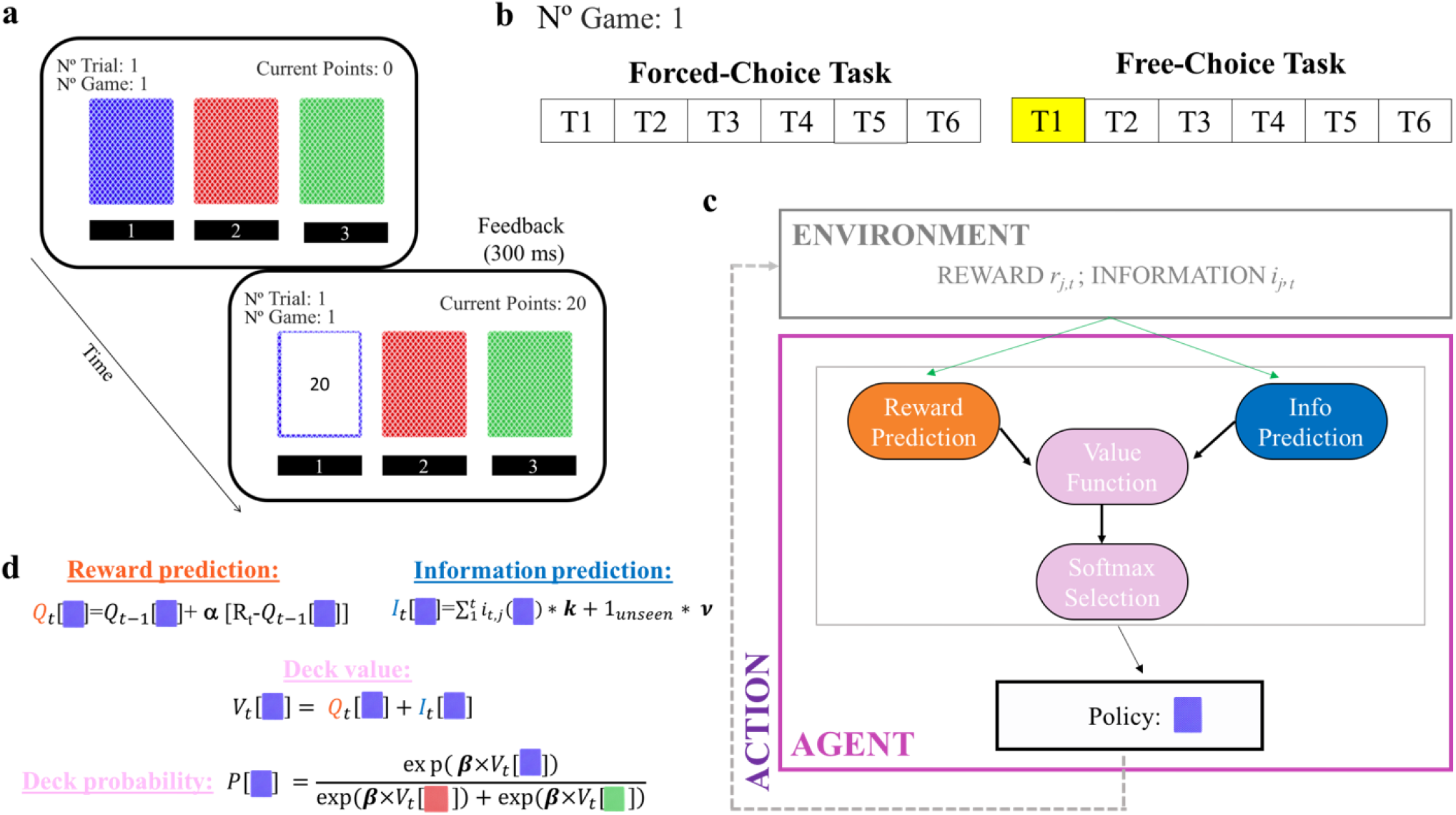
Behavioral task and RL model. **a)** On each trial, participants made choices among three decks of cards. After selecting a deck, the card flipped and revealed the points earned, between 1 and 100 points. Participants were instructed to attempt to maximize the total points earned at the end of the experiment. **b**) On each game, participants play a forced-choice task (6 consecutive trials) followed by a free-choice task (variable between 1 and 6 trials) on the same three decks. Subjects earned points only in the free-choice task. **c**) On each trial, the novelty-knowledge RL (nkRL) model computes an option value function according to both experienced reward and information associated with each option, then the model generates a choice by passing the option values through a softmax function. **d**) For each chosen option, nkRL uses a delta rule to update the reward prediction (α parameterized the learning rate), and updates information prediction as sum of general information (total number of times an option has been chosen) and a novelty term. The general information term describes the level of general information participants have about the selected option, while the novelty bonus is assigned to options the outcome of which has never been experienced in previous trials. Reward and information predictions are then combined into an overall action value, which are combined across options to through the softmax function (whose randomness is parameterized by the inverse-temperature parameter β). Model parameters are shown in bold.

When selected, each deck provided a reward (from 1 to 100 points) generated from a truncated Gaussian distribution with a fixed standard deviation of 8 points, and then rounded to the nearest integer. The generative mean for each deck was set to a base value of either 30 or 50 points and adjusted independently by +/-0, 4, 12, or 20 points (i.e., the generative means ranged from 10 to 70 points) with equal probability, to avoid the possibility that participants might be able to discern the generative mean for a deck after a single observation. The generative mean for each option was stable within a game, but varied across games. The generative mean reward value of the three decks had the same value in 50% of the games (*Equal Reward)* and with different values (*Unequal Reward*) in the other 50% of the games. In the Unequal Reward condition, the generative means differed so that two options had the same *higher* reward values compared to the third one in 25% of the games (*High Reward*), and in 75% of the games two options had the same *lower* reward values compared to the third one (*Low Reward*). The appearance of the reward conditions was randomized, as were the assignments of which two arms had the same generative mean within each game (in the Unequal Reward games).

On trials when participants do not choose the option with the highest reward expectation (or reward expectation is equalized across the choices), they can either choose at random (undirected or random exploration; ^4^, direct their exploration toward a novel option (novelty-seeking), or distribute their exploration among alternatives inversely proportional to how frequently they have been seen in the past (general information-seeking). In order to dissociate among these factors, we implemented two conditions in the forced-choice task ^4^. Participants were either forced to choose each of the three decks 2 times (*Equal Information*), or to choose one deck 4 times, a second deck 2 times, and the third 0 time (*Unequal Information*). In the latter condition, the “0 time” deck is perceptually familiar to participants (the stimulus is presented at the beginning of the game) but its reward distribution is novel to participants. While only the “0 time” deck is completely novel, the “2 times” deck should be relatively more information-rich than the “4 times” to the participants. 50% of the games were assigned to the Unequal information condition. The order of card selection was randomized in both information conditions, as was the occurrence of the equal and unequal information conditions.

### Computational model

We assume that humans behave according to both reward- and information-related internal beliefs/motivation when performing the above decision-making task ^6^. We formalize this using a reinforcement-learning (RL) type computational model (**Figure 1c**). In order to investigate the nature of information valuation in HCs and PGs, we implement a novel computational model that we term the “novelty-knowledge RL” (nkRL) model. NkRL combines reward and information evaluation using a delta learning rule ^30^ Eq. S1; **Figure 1d**), as in a previously proposed variant (Eq. S3; ^6^, but nkRL specifically dissociates the values associated with novelty and general information:

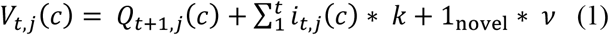

where *Q*_*t,j*_(*c*) is the expected reward value on trial *t* in game *j* for choice *c* (computed using Eq. S1), 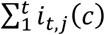 is the cumulative information about *c* acquired through trial t (*i*_*t,j*_ is 1 if selected on trial *t*, or 0 otherwise), and *k* is the *knowledge* (or general information) parameter which defines the weight toward previously acquired information (*k* being negative means there is a bonus toward lesser known options, while being positive means there is a bonus toward more familiar options). 1_novel_ ∗ *v* captures the value associated with *novelty*, where 1_novel_ is a Kronecker delta function that evaluates to 1 when *c* has never been seen in the current game and 0 otherwise, and the parameter *v* quantifies the value associated with novelty. As in previous algorithms in artificial intelligence, the novelty bonus is incorporated as optimistic initialization to the starting value of novel options ^31^. Lastly, we assume a choice is made via a softmax function of *V*_*t,j*_(c) ^32^ Eq. S2), where the decision policy is controlled by the inverse temperature β (**Figure 1d**). nkRL can shed light on the processes that underpin information valuation in both HCs and PGs by distinguishing the effects of reward-seeking and information-seeking on choices (β vs. *k, v*), and of novelty and general information on information-seeking (*v* vs. *k*). The model’s parameters are estimated by fitting nkRL to trial-by-trial participants’ free choices (**Supplement**).

## RESULTS

### Model-Free Results

#### Novelty-seeking in HCs and novelty-failure in PGs

We first examined how HCs and PGs compare in the influence of reward and information on choice behavior. We focus on the Unequal Information condition (equal information games have no informative options) and the first free-choice trial, the one trial where we can be sure that information and experienced reward are uncorrelated ^4^. We consider a trial to be *novelty-seeking* if the participant selects the novel option, and *reward-seeking* if the participant selects a previously experienced option with the higher empirical mean (regardless of whether it was seen twice or four times). For each subject, we computed the relative frequency of novelty-seeking trials and of reward-seeking trials over the total number of novelty-seeking and reward-seeking trials. We then entered these values into a mixed effects logistic regression predicting choice type (novelty-seeking, reward-seeking) with group (PGs, HCs) and reward condition (Low Reward, High reward) and their interaction as fixed effects and subjects as random intercepts (1|Subject). This standard random intercept model had lower BIC (6076.5) compared to a full random coefficient model (with random intercepts and slopes; BIC = 6110.2). First, consistent with previous studies using the same experimental design on healthy subjects ^6 9^, we found a main effect of reward (beta coefficient = −0.824 ± 0.104 (SE), z = −7.90, *p* < 10^−3^), with novelty-seeking generally more common in the Low Reward condition. More interestingly, we found a significant fixed effect of group (beta coefficient = 0.643 ± 0.268 (SE), z = 2.4, *p* = 0.016), with PGs engaging in less novelty-seeking and more in reward-seeking behavior (**Figure 2a**). The interaction between group and reward condition was not significant (beta coefficient = −0.144 ± 0.132 (SE), z = −1.093, *p* = 0.274), suggesting that the two groups did not differ in the way the reward conditions affected choice behavior.

**Figure 2.**
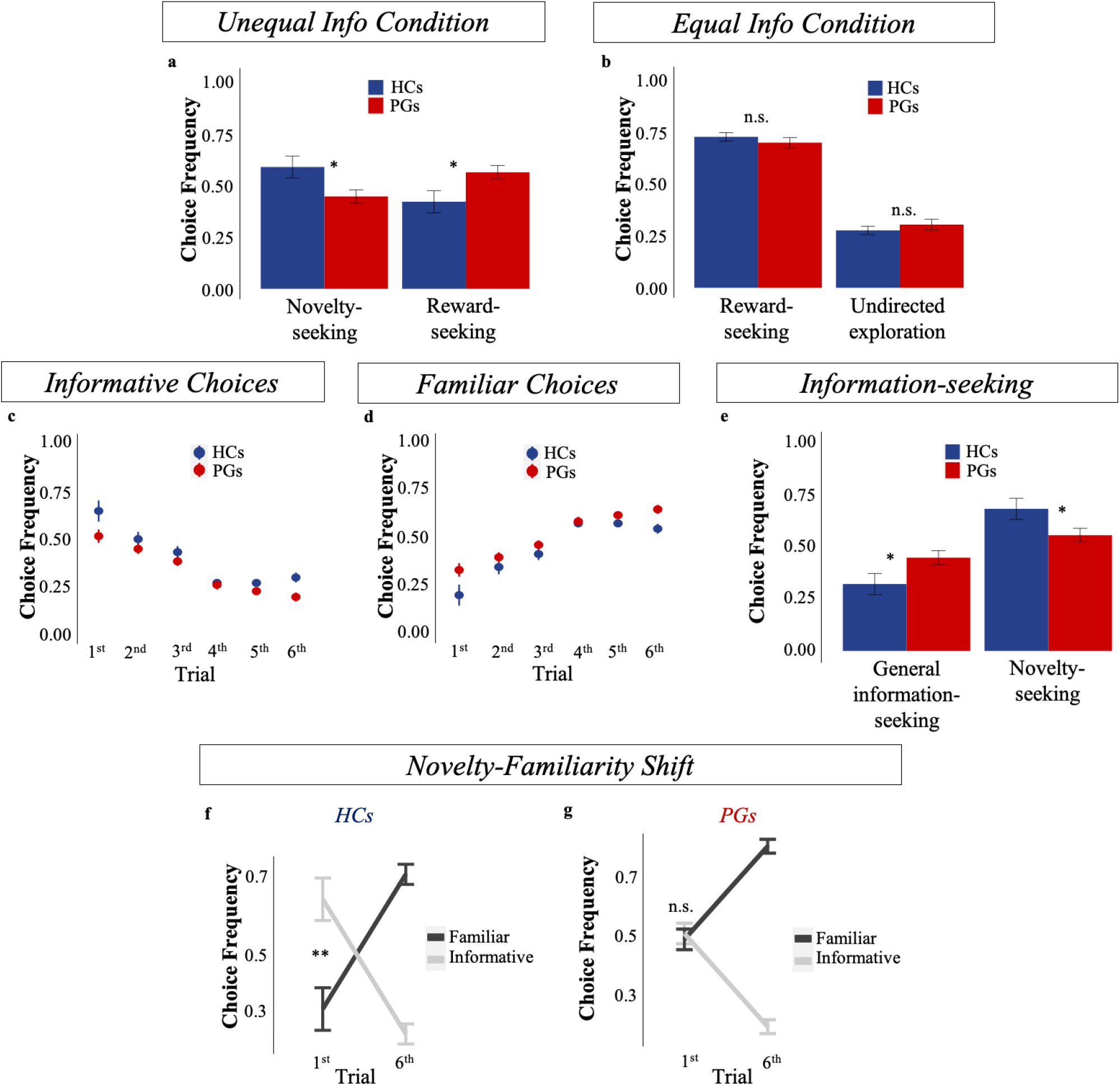
Model-Free analysis. **a**) Frequency of making *novelty-seeking* and *reward-seeking* choices over the total number of novelty-seeking and reward-seeking trials in the first free-choice trial of the Unequal Information condition (i.e., when options are sampled unequally during the forced-choice task; Unequal Info Condition in the figure). Novelty-seeking choices decreased and reward-seeking choices increased in PGs compared to HCs. **b**) Frequency of engaging in *reward-seeking* and *undirected exploration* in the first free-choice trial of the Equal Information condition (i.e., when options are sampled equally during the forced-choice task; Equal Info Condition in the figure). No difference was observed between the two groups. **c**) Frequency of selecting the option seen the *least* number of times in previous trial history (*informative choices*) in the Unequal Information condition. **d**) Frequency of selecting the option seen the *most* number of times in previous trial history (*familiar choices*) in the Unequal Information condition. In **c, d**, the frequencies were averaged across games in which participants were choosing informative and familiar options, thus the frequencies add to 1. **e**) Frequency of engaging in *novelty-seeking* and *general information-seeking* over the total number of information-seeking trials in the first free-choice trial of the Unequal Information condition: PGs have reduced information-seeking toward novel options (*novelty-seeking*), but increased information-seeking toward options selected twice in the forced-choice task (*general information-seeking*). **f**) HCs showed a novelty-familiarity shift: increased preference toward informative options in the first free-choice trial and an increased preference for familiar alternatives in the last free-choice trial. **g**) PGs showed no preference between informative and familiar options in the first free-choice trial, but a significant preference toward familiar options on the last free-choice. In all the figures, error bars represent standard error of the mean (s.e.m).

Interestingly, PGs and HCs show comparable choice behavior when choices were equally informative (Equal Information condition, **Supplement; Figure 2b**). This suggests that differences between the two groups were only present when choices were associated with different levels of information. Additionally, the shift in preference from more informative options (when subjects chose the option sampled the least number of times) early on in the free-choice task to more familiar options (when they chose the option sampled the most number of times) later on was smaller in PGs than HCs (**Supplement; Figure 2c,d**). Lastly, a “novelty-familiarity” shift was apparent in HCs (they prefer novel options in the first free choice trial) but absent in PGs who preferred novel options and familiar options equally on trial 1 (**Figure 2f,g**).

#### PGs have reduced preference for novelty but not for general information

The above analyses yielded hints that PGs have reduced preference specifically for novelty. To test this suggestion, we calculated the number of trials in which participants engaged in novelty-seeking and in general information-seeking (partially informative options sampled twice during the forced-choice task) and divided them by the total number of novel and general information trials to obtain their relative frequencies (i.e. we exclude trials in which the subject chose the option that was selected 4 times during the forced choice task). If alterations in PGs’ behavior are not specific to novelty, we should also expect to find lower selection of options experienced twice during the forced-choice task. Results showed that while PGs chose the novel option less often than HCs (*p* = 0.015; **Figure 2e**) on the first free-choice trial in the Unequal Information condition, PGs chose the partially informative option (seen twice) *more often* (M =0.446, SD = 0.21) compared to HCs (M =0.32, SD = 0.239; Wilcoxon Signed Rank test, *p* = 0.015; **Figure 2e**), suggesting that PGs specifically shy away from novelty-seeking and not from general information-seeking. As an additional check, we constructed a mixed logistic regression model to predict choice type (partially informative option, familiar option, i.e. excluding novel option trials) from group (PGs, HCs) as fixed effect and subjects as random intercept term (1|Subject; this model had lowest BIC compared to a model with random intercepts and slopes), and found no effect of group (beta coefficient = 0.011 ± 0.088 (SE), z = 0.12, *p* = 0.905), additionally suggesting no decrease in general information-seeking in PGs compared to HCs. We further examine this point in the next section.

### Model-Based Results

#### HCs have increased novelty bonus, while PGs have increased knowledge parameter

In order to elucidate the mechanisms underlying information-seeking in HCs and PGs, we turn to model-based analyses. Here, we propose a novel reinforcement learning-type model that we call “novelty-knowledge RL” (nkRL, **Methods**). We first ran a model comparison analysis (**Supplement**) and observed that nkRL was better able to explain participants’ behavior compared to the following models: a standard RL (sRL) model ^30^ - where only reward predictions influence choices; a knowledge RL (kRL) model ^6^ – which linearly combines reward and information associated with options without explicitly decomposing information into novelty and general information; leaky nkRL where information accumulation across trials proceeds in a leaky fashion; gamma nkRL (gnkRL) where information is measured sub- or super-linearly in the number of observations (**Figure 3a, b**; **Supplement**).

**Figure 3.**
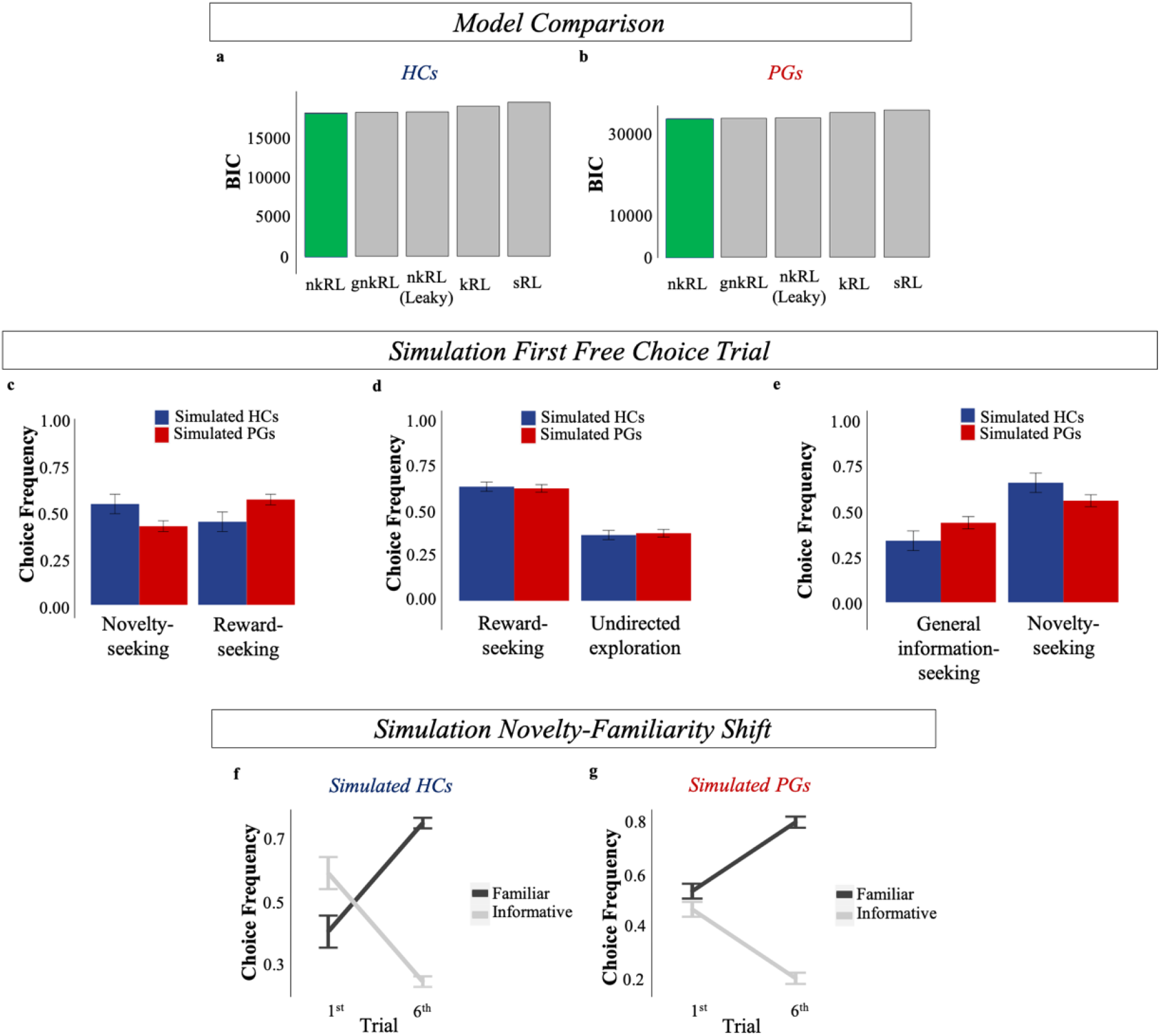
Model Comparison and nkRL simulations. BIC comparison of the 5 RL models in HCs (**a**) and PGs (**b**). The comparative fit is based on the sum of individual BIC computed by fitting each model to participants’ free choices. In both groups, novelty-knowledge RL model (nkRL, in green) better explains participants’ behavior compared to a leaky novelty-knowledge RL model (leaky nkRL), a knowledge RL model (kRL), a standard RL model (sRL) and a gamma novelty-knowledge RL model (gnkRL). By using the estimated individual parameters, simulations of nkRL in the first free choice trial reproduced the empirically observed decrease in novelty-seeking in PGs (Unequal Information condition, **c**), comparable choice behavior when choices are equally informative (Equal Information condition, **d**), an increase of preference for partially informative options (general information-seeking, **e**). **f**) nkRL correctly predicts the novelty-familiarity shift in the healthy sample, **g**) and its absence in the PG group. Error bars: s.e.m.

We then utilized nkRL to better investigate the process underlying the differences in information-seeking between PGs and HCs. We first simulated nkRL, using the individually fitted parameters, to verify that the model was able to replicate key behavioral patterns observed in the data. As shown in **Figure 3**, nkRL is able to qualitatively reproduce key behavioral patterns observed in both groups, including reduced novelty-seeking in PGs compared to HCs (**Figure 3c**), comparable choice behavior when choices are equally informative (**Figure 3d**), an increase of preference for partially informative options (general information-seeking, **Figure 3e**), and the absence of novelty-familiarity shift in PGs (**Figure 3g**).

Next, we performed parameter comparison analyses to examine which component of the decision-making process may be responsible for the behavioral pattern observed in PGs. We first performed a parameter recovery analysis to estimate the degree of accuracy of the fitting procedure (**Supplement; Figure S1**). We were able to recover all the parameters with high accuracy (all *r* > 0.8). We then compared the parameter estimates between the two groups. A Wilcoxon Signed Rank Test showed smaller novelty parameter *v* in PGs (M = 5.58, SD = 12.11) compared to HCs (M = 12.43, SD = 12.91, *p* = 0.0416; **Figure 4a**), while the knowledge parameter *k* was higher in PGs (M = 1.38, SD = 2.01) compared to HCs (M = 0.43, SD = 1.04, *p* = 0.0017; **Figure 4b**). In line with our model-free results, these results suggest that PGs have reduced information-seeking for novelty, but not for general information. We further explored this result by entering parameter (*v, k*) and group (HCs, PGs) in a two-way repeated measure ANOVA in a non-parametric setting using aligned rank transformation (e.g., ARTool package in R, http://depts.washington.edu/madlab/proj/art/;^33^. This revealed an effect of group (F(1,58) = 10.06, *p =* 0.002), an effect of parameters (F(1,58) = 40.19, p<10^−3^) and an interaction between group and parameter (F(1,58) = 18.13, *p* < 10^−3^). These results seem to confirm that the decrease in information-seeking in PGs is due to a failure in either computing or utilizing a novelty bonus and to increased importance to previously encountered but imperfectly explored alternatives. Interestingly, the two parameters interacted in a way that their relative difference was higher in HCs compared to PGs. To further investigate this, we computed the Euclidean distance between *v* and *k* (d_v-k_) in the parameter space. Results showed that d_v-k_ was larger in HCs (M = 14.9, SD = 9.3) than in PGs (M = 9.2, SD = 7.5) *p* = 0.034 (**Figure 4c)**. By simulating nkRL with low novelty parameters (i.e., small d_v-k_) and high novelty parameters (i.e., large d_v-k_), the model was able to predict the behavioral pattern observed in PGs and HCs, respectively (**Supplement; Figure S2**).

**Figure 4.**
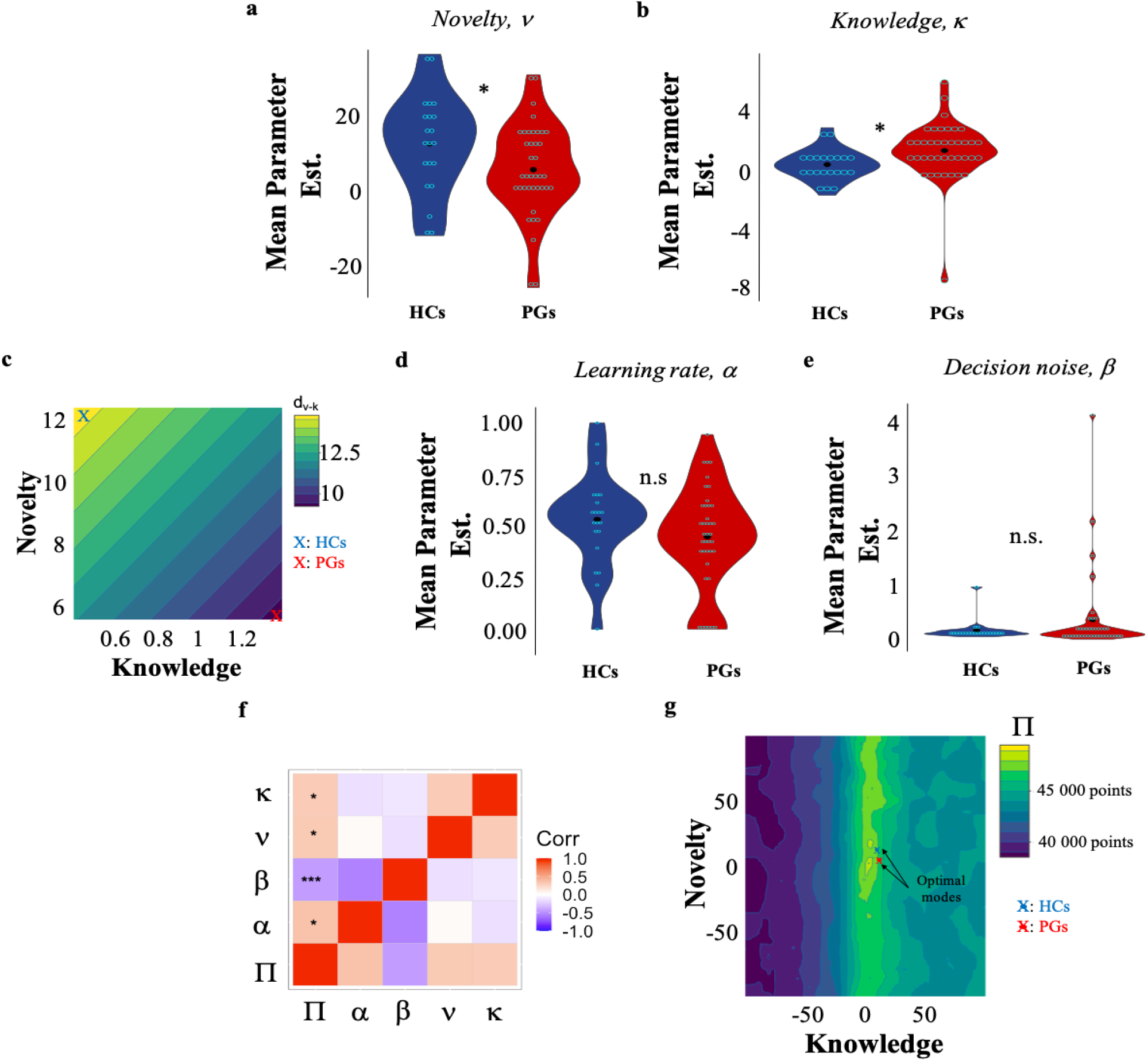
*nkRL* parameters and information-seeking modes. Model fit on all free-choice trials revealed a decrease in the novelty parameter ν (**a**) in PGs compared to HCs, while the knowledge parameter κ was higher in PGs compared to HCs (**b**). **c**) The Euclidean distance d_v-k_ between κ and ν was larger in HCs (in blue) than in PGs (in red). Learning rate α (**d**) and, decision noise β (**e**) did not differ between the two groups. **f**) Correlation matrix between nkRL model parameters and task performance П. P-values are corrected for multiple comparison (FDR). Both ν and κ positively correlated with П· Only correlations between П and model parameters are reported. **g**) Performance П across ν and κ parameter space. Averaged value of ν and κ for HCs is shown in blue, while in red for PGs. The two averaged values are expressed closer to the two optimal modes (in yellow).

Lastly, PGs and HCs did not differ in either learning rate α or softmax parameter β (p < 0.2; **Figure 4d**, **Figure 4e**) suggesting that the behavioral patterns observed in PGs were not related to learning alterations or due to an increase/decrease of random stochasticity in choice distribution. This latter result additionally confirms that exploratory impairments in PGs were specifically driven by novelty-related information valuation without affecting other undirected or unexplained exploratory components (e.g., softmax parameter). Overall, the model-based analyses appear to suggest that HCs are specifically driven by novelty during exploratory behavior (d_v-k_ is larger and in the direction of high novelty bonus; **Figure 4c)**, while in gamblers the integration of novelty is reduced and the integration of general information is enhanced resulting in a smaller distance between the two parameters.

#### HCs and PGs adopt distinct information-seeking modes

Previous analyses show that altered information-seeking in PGs compared to HCs was due to a decreased difference between *v* and *k*, with PGs given higher weight to knowledge and reduced weight to novelty. Here, we analyzed how this particular pattern might affect PGs’ reward accumulation performance in the task. We define task performance as sum of points earned on free-choice trials, summed across games. Our results showed no differences in task performance (П) between PGs and HCs throughout the task (all *p* > 0.05). We then correlated participants’ П with the estimated model parameters for each subject in both groups. We entered П and model parameters into a correlation matrix where p-values were corrected for multiple comparisons using False Discovery Rate correction (FDR ^34^. Results showed that both having increased novelty parameter and increased knowledge parameter relate to higher performance in the task (points earned; *p* < 0.05; **Figure 4f**). This seems to suggest that high novelty and high knowledge parameters are equally likely to yield high performance in our task. Additionally, no significant correlation was found between П and the distance d_v-k_ (p = 0.096).

The above results seem to suggest having a large (high novelty and low knowledge) or small (increased knowledge and decreased novelty) distance d_v-k_ yields good performance in the task. We further simulated the nkRL model with different settings of knowledge and novelty parameters, while keeping constant both alpha and beta parameters, to understand whether there were indeed two different modes that yield good performance in the task. We computed П for each simulation and we plotted it in the parameter space. Results showed that two modes gave high performance (**Figure 4g**): one mode with high novelty and low knowledge parameters (ν = 19.02; κ = 5.37, П = 48835 points) and a second mode with similar values for knowledge and novelty parameters (ν = 2.55; κ = 2.97, П = 49251 points). Interesting, average estimated values of ν and κ for the two groups were close to the two locally optimal modes. These results not only suggest that differences between HCs and PGs’ information-seeking behavior correspond to adopting two alternative modes of adaptive behavior for the task, but that reward feedback from the task would not be effective for shifting either group’s behavior to the alternative local optimum.

## DISCUSSION

In this study, we adopted behavioral, self-reported, and computational measures to investigate the underlying motivations driving information-seeking in healthy and addicted individuals. We focus on gambling disorder, a form of addiction without the confound of substance consumption ^35^ and characterized by compulsive gambling ^25^. We found that HCs and PGs adopt distinct information-seeking modes, closely related to the two locally optimal modes which yield to good task performance. HCs’ information-seeking behavior appears mostly driven by novelty-seeking with little effect of general information-seeking. On the contrary, PGs exhibit enhanced general information-seeking and reduced novelty-seeking compared to HCs. Our findings not only shed new light on the motivations driving healthy and addicted individuals to seek out information, but they also have important implications for treatment and diagnosis of behavioral addiction.

One possible interpretation of our results in regard to the difference between PGs and HCs is that novelty-seeking may be particularly important for human wellbeing and mental health, and the relative reduction of novelty-seeking may underlie the pattern of maladaptive behavior in problem gamblers compared to healthy controls. The link between novelty-seeking and wellbeing has been already suggested in previous research ^36^. More research is however needed to better understand the specific role novelty-seeking plays in human wellbeing and mental health. Nevertheless, novelty-seeking seems to have an adaptive role as it is expressed both in animal models and it motivates exploration in artificial agents. For example, animals can learn to press a key only for the sake of poking the head into a new compartment ^37^ or to guarantee the delivery of novel visual stimuli ^38^. To encourage exploration in artificial agents a fictive reward bonus is given to novel options ^39 31^, and similar heuristics seem to be adopted by the human brain ^12^. Novelty biases therefore might be crucial for quickly understanding changing environments, as when novel options are available for selection. The increased general information-seeking in PGs might be a compensatory mechanism which arises in addictive behaviors. To perform as healthy subjects, PGs increase general information-seeking as this also drives exploration of novel options (to a certain extent), while also enhancing additional exploration of other imperfectly known options in comparison to the healthy brain. It is worth noting that while both modes of behavior achieve good performance in our experimental task, in real life one may prove more maladapted than the other in many situations. Future work is needed to investigate this issue further.

Another possible interpretation of our finding is that there is a single underlying pattern of alteration in the brain structure of PGs that both reduces novelty-seeking and increases general information-seeking. Information-seeking behaviors are controlled by an interconnected cortico-basal ganglia network ^40^. Previous studies ^18 11 12 19 20 21 22^ seem to support the dissociation of these two motivational factors within this network. Further work however is needed to individuate how the neural markers for novelty and general information interact within the information-seeking network and can produce the altered behavioral pattern observed in PGs.

An interesting implication of our findings is that, regardless of the provenance of the alternative pattern of information-seeking in PGs compared to HCs, this apparent reduction of novelty-seeking and increase in general information-seeking may be useful for developing novel diagnostic tools and even novel treatments for this pathology. First of all, our findings potentially suggest a novel method for identifying individuals with behavioral addiction i.e. reduced novelty drive and increased general information seeking. Second of all, novel theories on the pathophysiology of this disorder suggest that the resolution of reward uncertainty present in gambling games creates the capacity for addiction ^27 41 26^. Our findings can help clarify why addictive behaviors are characterized by reduced information-seeking ^10^, and yet the source of addiction involves resolving uncertainty. Interestingly, reward uncertainty in addictive behaviors hijacks the dopaminergic system ^27 41 42^, as drugs do in substance addiction. Given that novelty-seeking relies on the functioning of midbrain dopaminergic system ^43 12 44^, this behavior may compete with responses towards reward uncertainty. In other words, reduced novelty-seeking might be a signature of this hijacking process.

Our study however does not rule out the possibility that neurophysiological alterations in the brain could pre-date or even induce problem gambling. In particular, it might be possible that individuals who show the “reduced novelty-seeking and increased general information-seeking” mode may be more predisposed for developing addiction. When addictive behaviors arise, the reduced ability to represent novel behavioral patterns may freeze their decision processes and trap them into the same behavioral routines. Reduced novelty-seeking might therefore explain why addicted individuals are trapped in the same behavioral routines despite the negative consequences associated with them (e.g., financial loss^25^.

On additional note, novelty seems to compete with conditioned drug rewards ^45^. This may suggest that boosting novelty-seeking behavior may compete with the addictive stimuli and reduce the impact of addiction. Therefore, novelty-seeking might be introduced in current treatments for addictive behaviors ^46 47^. Further work however is needed to test whether our findings can be extended to effective clinical interventions. Additionally, while reduced information-seeking has been observed in both behavioral ^10^ and substance addiction ^24^, our study cannot inform on whether this dissociation is a common code for addiction or instead it is only key to behavioral addiction. Further work is needed to test whether our findings can be generalized to other addiction types.

Concerning healthy information-seeking, our results show a more nuanced view over information-seeking under repeated choices (or directed exploration; Wilson et al., 2014). While in previous RL models directed exploration was modelled as general information or uncertainty parameter added to the value function ^4,48^; ^6 49 8^, here we were able to dissociate the contribution of novelty-seeking and general information-seeking to human exploration. We observe that a novelty bonus and general information can play dissociable roles, with potentially different implications for different decision-making scenarios or exploratory phases. Our findings therefore strengthen the view of exploration as a multifaceted and sophisticated process ^4 49^. Moreover, our results replicate previous findings that assign different behavioral roles and neurocognitive mechanisms to informative and undirected components of exploration ^4,6,9,50-52 53^. Indeed, we found PGs display reduced directed exploration (defined here as choosing the most informative option- the novel option) but not undirected (or random) exploration (both in terms of softmax parameter and exploratory choices made in the Equal Information condition).

Altogether, our findings extend the scientific understanding of human information-seeking in healthy and addictive behaviors. HCs and PGs showed distinct information-seeking modes. Healthy information-seeking was motivated by novelty, while PGs’ information-seeking was characterized by reduced novelty and increased general information. Our results suggest that the expression of novelty-seeking behaviors might be a potential predictor of human wellbeing, and the expression of altered information-seeking patterns a potential marker of addiction. Methodologically, this work offers promising novel experimental and computational approaches for studying the mechanisms underlying information-seeking under repeated choices in both healthy and pathological populations.

## ACKNOWLEDGEMENT AND DISCLOSURE

ICD was supported by F.R.S.-FNRS grant (Belgium). XN is a research associate the F.R.S.-FNRS (Belgium). AC is a research director at the F.R.S.-FNRS (Belgium). AJY is funded in part by an NIH/NIDA CRCNS grant (R01 DA050373-01). This work was in part funded by ERC Advanced Grant RADICAL to AC. We would like to thank Pauline Deroubaix for helping I.C.D in recruiting participants and collecting their data and Tali Sharot for comments on previous versions of this manuscript. The authors report no competing financial interests.

## SUPPLEMENTARY MATERIAL

### SUPPLEMENTARY METHODS

#### Clinical and demographic characteristics

Inclusion/exclusion criteria were examined the day before the experiment by conducting a short telephone interview as well as on the day of the experiment by filling self-reported questionnaires presented in a random order during the last part of the experimental session. The telephone interview was adopted as pre-screening for both PGs and HCs. We specifically asked for information concerning age, gender, frequency of gambling per week (for PGs) or last gambling experience (for HCs), consumption of alcohol per week or substance (including legal and illegal drugs), inability to stop drinking alcohol, undergoing psychological treatments, and possible brain surgeries underwent in the past. We interviewed about N=60 gamblers. Gamblers who met the criteria were then invited to take part to the experiment (N=40). We then took the demographics of the gambling group (gender and age) and we set them as criteria for selecting the control group (alongside with no gambling experience in the year before the study, no sign of excessive use of alcohol or use of substances, psychological treatments, possible brain surgeries etc.). We interviewed about the same number of participants as for the gambling group. More than half of the sample was rejected because of gender (as the gambling group was mostly composed of males) and age (gamblers were quite old compared to usual undergraduates or master students who take part to psychological experiments at the University). In the following two sections, we describe the clinical and demographic characteristics of PGs and HCs.

##### Problem gamblers

Gambling severity was evaluated using the Canadian Problem Gambling Index (CPGI ^1^). Eight gamblers were classified as low level of problem gambling with 1≤ GPCI ≤ 3, thirteen gamblers with moderate level of problem gambling (leading to some negative consequences; 4 ≤ GPCI ≤ 7), and nineteen as exhibiting pathological problem gambling (with negative consequences and possible loss of control; GPCI≥8). We also interviewed participants using DSM-V (French translation) and we observed that 52.4% of PGs met the DSM-V criteria for gambling disorder ^2^. The relatively low level of gambling addiction presented in this population is the result of including only participants who showed no co-morbidities with substance abuse or alcohol use disorder. Specifically, to be able to tell apart effects of addictive behaviors *per se* on decision-making from effects of long-term intake of chemical compound, we tested PGs with no use (N= 31, Drug Abuse Screening Test ^3^- DAST =0) or non-problematic use (N=9, DAST =1) of legal and illegal substances and with absence of alcohol addiction (Alcohol Use Disorders Identification Test ^4^- AUDIT-<12 in men and AUDIT < 11 in women, M = 4.625, SD = 3.868; N=30 did not show any misuse of alcohol AUDIT< 8). We also controlled for smoking addiction using the Fagerström Test for Nicotine Dependence-FTND ^5^. Seven participants reported to smoke, but only 2 were classified with a mid-dependence and 2 with a weak-dependence, the other 3 were not dependent. Given that the main statistical results remained unchanged after removing those participants, we decided to include them in all the analyses. Additionally, to avoid the scenario that participants under psychological treatment may have developed a certain type of cognitive strategy over their decision processes, we included only participants who were not undergoing or seeking for psychological treatment. Moreover, we only included regular gamblers that were gambling at least once per week. Finally, we recruited both strategic PGs (sport betting, poker, black jack; N=22) and non-strategic PGs (bingo, lotto, slot machine, roulette; N=18) ^6^. Given that no behavioral difference was found between the two sub-types (in line with ^7^), we combined strategic and non-strategic gamblers in the same gambling group in all analyses reported in this manuscript.

##### Healthy controls

The inclusion criteria for the HC group were as follow: CPGI=0 and no gambling experience in the past 12 months. 40% of control participants reported to have gambled in the past years, whereas the rest of the group reported to have never gambled in their life. As for the problem gambling group, we only included participants who scored DAST < 2 (with 17 subjects DAST = 0) and AUDIT < 12 (for the men), 11 (for the women) (with 17 subjects scored AUDIT< 8; M = 5.3, SD = 3.1). Three participants reported to smoke, two of them showed no sign of addiction (FTND = 0; 2) and one showed mid-level of addiction (FTND = 7). Removing this participant did not change the main statistical results, therefore the participant was included in all the analyses.

#### Behavioral Task

To study information-seeking behavior under repeated choices, we adopt a modified version of a popular task (i.e., the multi-armed bandit) often used to study sequential learning and decision-making behavior. In the bandit task, the decision-maker must make repeated choices among options characterized by initially unknown reward distributions. Each choice can be driven either by a more myopic desire to maximize immediate gain (based on knowledge gained from previous choices and outcomes) or by a more long-term goal of being more informed about all the options. In these repeated scenarios, however, the more the decision-maker tends to choose the most rewarding options, the more those rewarding options tend to be (anti-) correlated with the amount of (remaining) information that can be obtained ^8 9^. Accordingly, these classical decision-making tasks make it difficult to quantify exactly how much reward and information each contribute independently to choices ^9^. Here, we therefore adopt a novel variant of the bandit task ^10^, inspired by ^9^, which has an initial phase of forced choices that carefully controls for reward and information associated with each option. In particular, the influence of reward and information on choices is orthogonalized in the first free-choice trial (since after receiving the feedback on the first free-choice trial, subjects tend to choose the more rewarding options more often, thus reward and information become anti-correlated). Adding a forced-choice task before the actual decision task allows to control for available information and the reward magnitude associated with each option (i.e., options associated with the lowest amount of information were least associated with experienced reward values)^9^. This procedure allows to dissociate between information-driven exploration and undirected exploration. For instance, in the unequal sampling condition, the deck never selected during the forced choice task has highest informative value (it is completely unknown to participants) but it has no reward value associated with. By choosing that deck, participants are engaging in information-driven exploration. On the contrary, in the equal information condition, no differences are observed in terms of information. Therefore, whenever participants choose to explore, this strategy is not driven by an informative drive but only by decision noise^9^.

Considering only the first free-choice trial (the trial where reward and information are least correlated ^9^), we then define three types of behaviors, corresponding to three distinct motivational factors: (1) *Novelty-seeking exploration* refers to choosing the novel, never-seen option in the Unequal Information condition; (2) *General information-seeking* refers to choosing partially informative options sampled twice in the Unequal Information condition - these options are still informative when explored but not completely novel; (3) *Reward-seeking* refers to choosing options associated with the highest gain. Additionally, we define a fourth behavior - *undirected exploration-*which refers to choosing options associated with the lowest gain in the Equal Information condition, as this type of choice is neither driven by reward nor by information-seeking.

Contrary to our previous versions of this task ^10 11^, in half of the games of the equal reward-equal information condition, we introduced an unusually high reward outcome (with respect of the deck mean in that game) for a specific option (e.g., 90 points) the first time that this option was selected in the forced-choice task (subsequently the mean of the deck was set to its original value). This manipulation was introduced as a control condition in order to test whether gamblers’ perseverate in choosing a generally poor option that they initially have a good experience with (the ‘big win’ hypothesis for gambling addiction ^12^).

Prior to beginning the main experiment, participants were told that during the forced-choice task, they may sample options at different frequencies, and that the decks of cards did not change during each game, but were replaced by new decks at the beginning of each new game. However, they were not informed of the details of the reward manipulation or the underlying generative distribution adopted during the experiment.

#### Computational Modelling

In this section, we provide details on the RL models adopted in this study.

##### Standard RL model

The standard RL (sRL) model learns reward values on each trial using the delta learning rule^13^

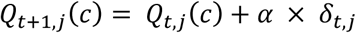

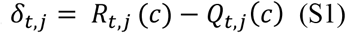

where *Q*_*t,j*_(*c*) is the expected reward value for trial *t* and game *j* and *δ*_*t,j*_ is the *prediction error*, which quantifies the discrepancy between the previous predicted outcome *Q*_*t,j*_(*c*) and the actual outcome *R*_*t,j*_ obtained at trial *t* and game *j*. Since participants were told that games were independent from one another, *Q*_0_ is initialized at the beginning of each game to the global estimate of the expected reward values for each deck. We previously showed that this initialization was better able to capture healthy participants’ behaviour than learning *Q*_0_ on a trial-by-trial basis ^10^. Next, a choice is made by entering expected reward values into the softmax function ^14^, as follows:

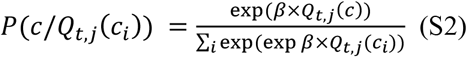

where *β* is the inverse temperature that determines the degree to which choices are randomized by decision stochasticity (or choice variability).

##### Knowledge RL model

As sRL, the knowledge RL (kRL) model learns reward values using Eq. S1 but it additionally integrates information obtained from each deck into the value function:

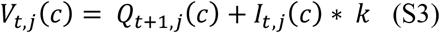

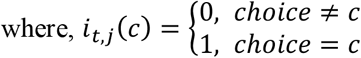

κ modulates the importance of information relative to experienced reward. With large κ the model favors already experience decks, while with negative values of κ the model explores new information more frequently. A choice is made by entering choice values *V*_*t,j*_(*c*) into Eq. S2.

##### Novelty-knowledge RL model

As the above models, the novelty-knowledge RL (*nkRL*) model learns reward values using Eq. S1. And, it additionally integrates information into the value function as kRL. However, as described in the main text, nkRL computes information as a sum of knowledge term and novelty term resulting in the following value function:

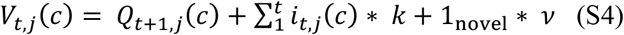

A choice is made by entering choice values *V*_*t,j*_(*c*) into Eq. S2.

##### Leaky nkRL model

The leaky nkRL model learns reward values using Eq. S1 and it integrates both knowledge and novelty term into the value function as nkRL. However, in leaky nkRL each bit of new information is integrated in a leaky fashion as follow:

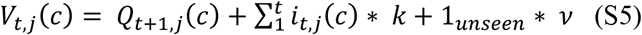

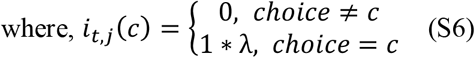

##### Gamma nkRL model

The gamma nkRL (gnkRL) model learns reward values using Eq. S1, and it integrates both knowledge and novelty term into the value function as nkRL. However, gnkRL allows a non-linear integration of information:

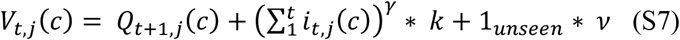

γ defines both the degree of non-linearity in the amount of observations obtained from options after each observation and its related importance. Under high γ the information already gained is highly relevant, whereas the information to be acquired is less relevant or penalized. γ is constrained to be > 0.

##### Model fitting and Model selection

The models’ parameters were estimated by fitting the model to trial-by-trial participants’ free choices (∼600 choices for each subject). The fitting procedure was performed using MATLAB function *fminsearchbnd* and iterated for 15 randomly chosen multiple starting points in order to minimize the chance of finding a local optimum instead of a global one. The fitting procedure was validated by running a recovery analysis: the model was simulated on the task using the retrieved parameter estimates to generate synthetic behavioral data and then the fitting procedure was applied to the synthetic data in order to check whether previously estimated parameters were indeed recovered ^15^ (**Figure S1**). For model comparisons, negative log likelihoods obtained during the fitting procedure were used to compute model evidence (the probability of obtaining the observed data given a particular model). We adopted an approximation to the (log) model evidence, namely the Bayesian Information Criterion (BIC) ^16^ and we compared its estimate across different models (fixed-effect comparison). Additionally, we used random-effects procedure to perform Bayesian model selectin at group level ^17^. In order to inspect the fitting procedure for overfitting we adopted cross validation procedure ^18^. We fitted the model to 70% of the trials and we tested its ability to predict choices on future data (30% of the trials) compared to a simpler nested model. We then adopted the likelihood ratio test to determine if the better fit of complex model was due to noise captured in the data.

#### Statistical analysis

Statistical analysis was performed using RStudio (https://www.rstudio.com/). When violations of parametric tests were indicated, non-parametric tests were performed. *P*-values < .05 were considered significant.

### SUPPLEMENTARY RESULTS

#### PGs and HCs show comparable choice behavior when choices are equally informative

The reduced novelty-seeking behavior in PGs found above could either be due to a specific decrease in the valuation of novelty, or a relative and general increase in the valuation of reward. To examine this, we compare the two groups’ behaviors in the Equal Information condition, in which the options have been sampled equal number of times and thus equally informative – any systematic difference in reward-seeking behavior here would be attributable specifically to reward and not influenced by general information or novelty. Again, we focus on the first free-choice trial, where there is no confound between reward and information. We classified choices as *reward-seeking* when choosing the option associated with the highest amount of points and *undirected exploration* otherwise. We then entered these values into a mixed effects logistic regression predicting choice type (reward-seeking, undirected exploration) with group (PGs, HCs) and reward condition (Low Reward, High Reward) and their interaction as fixed effects and subjects as random intercepts (1|Subject). This model had lower BIC (5956.2) compared to a model with random intercepts and slopes (BIC = 5977.7). Replicating previous studies using the same experimental design on healthy participants ^10 11^, we found a fixed effect of reward (beta coefficient = −0.351 ± 0.109 (SE), z = - 3.23, *p* < 10^−2^), with undirected exploration lower in the Low Reward condition. The fact that low reward enhances novelty-seeking but reduces undirected exploration suggests that these are dissociable exploratory drives in the brain with dissociable neural substrate ^11,19^. Most importantly, the effect of group (beta coefficient = 0.113 ± 0.191 (SE), z = 0.589, *p* = 0.556) and the interaction between group and reward (beta coefficient = −0.016 ± 0.135 (SE), z = 0.116, *p* = 0.908) were not significant. The results from the current analysis, along with those from the previous analysis, suggest that the reduced novelty-seeking behavior in PGs is specific to novelty and not an indirect consequence of greater valuation of immediate reward in general (**Figure 2b**).

#### Novelty-familiarity shift is absent in PGs

Here, we examine choices made by participants across the entire set of free choice trials in the Unequal Information conditions. We classified a choice as an *informative choice* when subjects chose the option sampled the least number of times thus far, and *familiar choice* when they chose the option sampled the most number of times so far. We calculated the number of trials in which each choice was made and divided them by the total number of informative and familiar trials to obtain their relative frequencies (i.e. we exclude trials in which the subject chose the option that was neither most familiar nor most informative). We then entered those values into a mixed effects logistic regression predicting choices (informative, familiar) from group (PGs, HCs) and trial (1,2,3,4,5,6), and their interaction as fixed effects and subjects as random intercepts (1|Subject; this model had lower BIC compared to a model with random intercepts and slopes). This revealed a fixed effect of group (beta coefficient = 0.546 ± 0.203 (SE), z = 2.69, *p* = 0.007) and fixed effect of trials (beta coefficient = 0.419 ± 0.021 (SE), z = 19.58, *p* < 10^−3^) as when both equal information and unequal information games were included (**Figure 2c, d**). However, narrowing the analysis to the Unequal Information condition also revealed an interaction effect between group and trial (beta coefficient = −0.06 ± 0.027 (SE), z =-2.23, *p* = 0.026), such that the shift in preference from more informative options early on in the free-choice task to more familiar options later on was smaller in PGs than HCs. To better understand this interaction, we compared subjects’ tendency to choose the most informative versus most familiar option on the first and sixth trial of the free choice task. We found that control subjects preferred novel options (M= 0.641, SD= 0.257) over familiar options (M= 0.359, SD= 0.257; *p* = 0.002; **Figure 2f**) on trial 1, but reversed preferences to prefer familiar options over informative options on trial 6 (M = 0.705, SD = 0.121, *p* < 10^−3^). In contrast, PGs preferred novel options (M= 0.51, SD = 0.222) and familiar options (M= 0.49, SD = 0.222) equally on trial 1, but strongly preferred familiar options (M= 0.807, SD = 0.149, *p* < 10^−3^) over informative options (M= 0.193, SD = 0.149) on trial 6. Thus, the “novelty-familiarity” shift was apparent in HCs but absent in PGs.

#### Model comparison

We first examine whether our nkRL model was better able to explain participants’ behavior compared to a standard RL (sRL) model ^13^ -where only reward predictions influence choices-and, to a knowledge RL (kRL) model ^10^ –which combines both reward and knowledge associated with options without explicitly decomposing information into novelty and general information. We chose kRL as example of unitary models (i.e., information is not decomposed in different drives) because previous researches showed that kRL was better able to explain human behavior in our behavioral task compared to models which update learning rate as number of observations (e.g., Kalman filter, ^10^). We fit the 4 models to participants’ data and we computed model evidence as approximation of –BIC/2. We removed two subjects (one from each group) for bad fitting. These subjects were removed from all model-based analyses reported in the main text. We then utilized Bayesian Model Selection ^17^ to compare the 3 models. We found nkRL model was the best model for predicting choice behavior in both HCs (xp_nkRL_=1, BIC_nkRL_=18065.6; xp_kRL_=0, BIC_kRL_= 18918; xp_sRL_=0, BIC_sRL_= 19407; **Figure 3a**) and PGs (xp_nkRL_=0.877, BIC_nkRL_= 33577.2; xp_kRL_=0.058, BIC_kRL_= 35080.1; xp_sRL_=0.065, BIC_sRL_= 35683.9; **Figure 3b**). Next, we asked whether participants were integrating complete information into the value function, as predicted by nkRL, or instead information was integrated in a leaky fashion. We implemented a new model (leaky nkRL) where each sample of information integrates as 1*λ, where λ is the leaky integration parameter. Model comparison showed that nkRL model was better able to explain both PGs (xp_nkRL_= 0.9999, BIC_nkRL_= 33577.2; xp_leaky_nkRL_=0.0001, BIC _leaky_nkRL_=33795.2; **Figure 3b**) and HCs’ choices (xp_nkRL_= 1, BIC_nkRL_=18065.6; xp _leaky_nkRL_=0, BIC _leaky_nkRL_=18188.7; **Figure 3a**). Lastly, we examined how information affects choice values. It may be the case that at least for certain situations (as in the present task) in which only a few samples of each option are available, additional observations may provide a non-constant amount of information and therefore they may scale choice value in a sub or super-linearly fashion. We compared nkRL, where information is measured linearly in the number of observations, with a model that permits the integration of information sub- or super-linearly (gnkRL). Model comparison showed that nkRL model was better able to explain both PGs (xp_nkRL_=1, BIC_nkRL_= 33577.2; xp_gnkRL_=0, BIC_gnkRL_= 33703.9; **Figure 3b**) and HCs’ choices (xp_nkRL_=1, BIC_nkRL_= 18065.6; xp_gnkRL_=0, BIC_gnkRL_= 18137.4; **Figure 3a**). Thus, we found nkRL to be the best-fitting model among all those that we considered.

#### Parameter recovery

We performed a parameter recovery analysis to estimate the degree of accuracy of the fitting procedure. To do so, we simulated data from nkRL using the parameters obtained from the fitting procedure (*true parameters*), and we fit the model to those simulated data to obtain the estimated parameters (*fit parameters*). We then ran a correlation for each pair of parameters ^15^ (**Figure S1**). This revealed high correlation coefficients for alpha (r_HCs_ = 0.8, *p*_HCs_ < 10^−3^; r_PGs_ = 0.9, *p*_PGs_ < 10^−3^), knowledge (r_HCs_ = 0.9, *p*_HCs_ < 10^−3^; r_PGs_ = 0.6, *p*_PGs_ < 10^−3^) and novelty (r_HCs_= 0.98, *p*_HCs_ < 10^−3^; r_PGs_ = 0.8, *p*_PGs_ < 10^−3^). The beta parameter showed high correlation coefficient in PGs (r = 0.9, *p* < 10^−3^). In HCs one participant showed bad fitting while the rest of the group showed high correlation coefficient (r = 0.97, *p* < 10^−3^). We removed this participant during the comparison of the beta parameter.

#### Simulations nkRL with random parameters

In this section, we report the result of the simulation of the nkRL model with random parameters to better understand the effect of novelty on choice behavior. We simulated nkRL with High Novelty and Low Novelty parameter. In each set of simulations, nkRL was simulated 100 times. In High Novelty, the averaged values of the parameters were as follow: alpha (M = 0.513, SD = 0.315), beta (M= 0.52, SD = 0.283), knowledge (M = 0.493, SD = 0.288), novelty (M = 41.38, SD = 11.31). In Low Novelty, we used the following averaged values: alpha (M = 0.519, SD = 0.304), beta (M = 0.51, SD = 0.293), knowledge (M = 0.479, SD = 0.282), novelty (M = −0.839 SD = 0.584). We then classified model choices in reward-seeking (when the model chooses the experienced decks with the highest average of points regardless of the number of times that deck had been selected during the forced-choice task) and novelty-seeking (when the model selects the option never sampled during the forced-choice task) in the first free-choice trial of the unequal information condition. As shown in **Figure S2a**, under Low Novelty the model increases reward-seeking at the expense of novelty-seeking as observed in PGs (**Figure 2a**). Next, we calculated the number of trials in which the model was choosing the partially informative option (seen twice) in the first free-choice trials of the unequal information condition and we averaged those estimates across the trials in which the model engages in information-seeking (novelty-seeking + general information-seeking). As shown in **Figure S2b**, under Low Novelty the model increases the selection of options selected twice during the forced-choice task (general information-seeking) at the expense of novel options as observed in PGs (**Figure 2e**).

#### Personality traits

In this section, we explore the individual differences between PGs and HCs to investigate whether personal traits could explain the behavioral differences observed throughout our analyses. We focus on intolerance of uncertainty (EII ^20^), impulsivity (UPPS-P ^21^), sensation-seeking (SSS ^22^), and sensitivity to punishment and reward (SPSRQ ^23^). Comparisons between HCs and PGs revealed no differences in the scores obtained from EII (*p* = .785, BF_01_ = 3.61), UPPS-P (*p* = .217, BF_01_ = 1.89), SSS (*p* = .483, BF_01_ = 3.02), and SPSRQ (sensitivity to reward *p* = .399, BF_01_ = 2.81; sensitivity to punishment *p* = .266, BF_01_ = 2.4), suggesting that the behavioral alterations observed in PGs are unlikely to be explained as differences in terms of personality traits (or in some cases there was not substantial evidence in favor of the alternative hypothesis). These results appear to suggest that reduced novelty-seeking in PGs may relate to a process or mechanism that is independent from individual subjective preferences toward uncertainty, sensation-seeking, or punishment and reward sensitivity.

#### The ‘big win’ hypothesis

The results reported in this study showed that PGs reduced novelty-seeking behaviors as a consequence of a failure to represent or incorporate a novelty bonus. However, these parametric alterations might have been confounded by the inability of PGs of moving away from an option after experiencing fairly positive outcomes in the past, i.e., the ‘big win’ hypothesis. To better investigate this point, we computed the empirical probability of choosing an option associated with an unusually high score (“big win” options) when first selected in the forced-choice task. A two-sample t test showed no differences in the probability of choosing the “big win” option in PGs (M = 0.607 SD = 0.187) compared to HCs (M = 0.596 SD = 0.144), *p* = .798 suggesting that PGs’ choice behavior was not driven by the persistence in choosing options associated with unusually good outcomes in the past.

## SUPPLEMENTARY FIGURES

**Figure S1.**
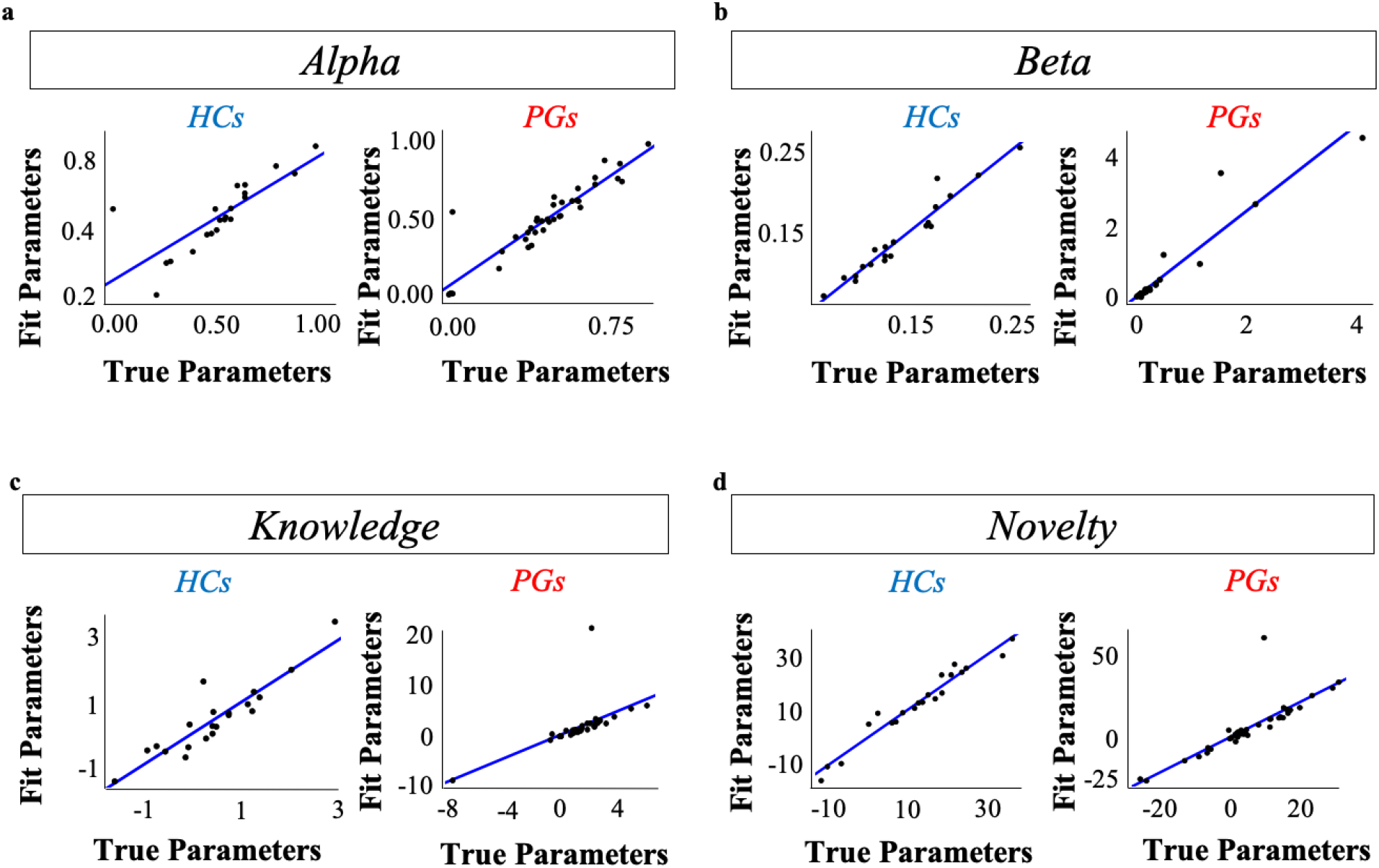
Parameter Recovery. Correlation between true and fit parameters for nkRL model. True parameters are those recovered during the fitting procedure, while fit parameters are those recovered after fitting the model to synthetic data (obtained by simulating nkRL with parameters estimated in the two groups).

**Figure S1.**
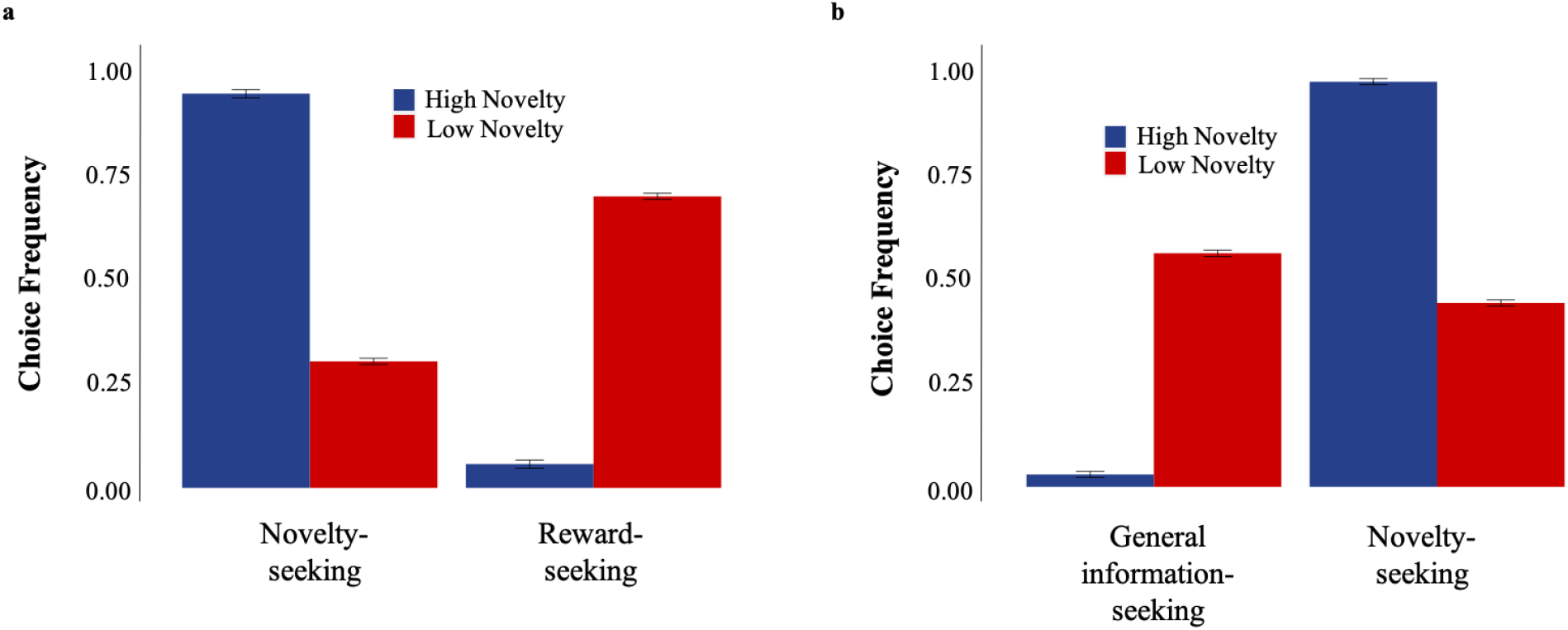
nkRL simulations with random parameters. Under Low Novelty the model frequently engages in reward-seeking (**a**) and in general information-seeking (**b**).

